# Vegetation and soil seedbank dynamics in *Parthenium hysterophorus* L. invaded subtropical grassland in Nepal

**DOI:** 10.1101/760561

**Authors:** Maan Bahadur Rokaya, Jyoti Khatri-Chettri, Shiba Raj Ghimire, Bharat Babu Shrestha

## Abstract

*Parthenium hysterophorus* is a noxious invasive weed and is ever expanding in its introduced range including Nepal. Understanding vegetation dynamics including soil seedbank in *Parthenium* invaded communities and the growth pattern of the weed itself is essential for effective management of *Parthenium*. We monitored growth of *Parthenium* (height, density, cover and soil seedbank) and plant species composition of associated species for 5-year period from 2009 in a grassland invaded by *Parthenium* in south-central Nepal. We found that *Parthenium* cover and height decreased from 2009 to 2010 and then slightly increased in 2013. *Parthenium* density decreased from 2009 to 2010 and then was variable until 2013. Year × grazing interactions had significant effect on *Parthenium* cover and density. *Parthenium* soil seedbank was eight times higher near the soil surface than in deep soil. It increased from 2009 to 2012 but decreased in 2013. Seedbank was also affected by interactions of year × depth, depth × grazing, and year × depth × grazing. Altogether, 87 plant species were recorded in *Parthenium* invaded sites and their species richness decreased until 2012 but slightly increased in 2013. The composition of associated plant species was affected by animal grazing intensity, *Parthenium* density, cover, and their interactions. *Parthenium* invasion has been ever increasing in our study site and many palatable plant species are under potential threat. Thus, there is an urgent need to carry out awareness campaign, formulate proper management plans, and implement such plans properly to manage *Parthenium* weed in Nepal.

## Introduction

In terrestrial ecosystems, impacts of biological invasions on associated plant community structure is observed high when the invader is producer and the recipient ecosystem is grassland [1]. This condition leads to the loss of native species and economic damages [2]. Invasiveness of an invader depends on persistence of seeds in the soil [3]. Once the status of seedbank is known it would be easy to formulate the control and management measures for invasive species. Though seedbank is difficult to destroy but reducing seed output could be achieved by using biological control agents [4] or destroying it mechanically, or by using chemicals [5]. In Nepal, seed bank of *Parthenium hysterophorus* L. (Asteraceae) (hereafter named as *Parthenium*) was investigated [6] but the long-term patterns are not well documented. Thus, we have investigated how the seedbank of *Parthenium* changes over a 5-year period in a same locality in Nepal.

Invasive species are known to have a wide range of ecological impacts to recipient ecosystems; these includes changes in nutrient cycling and soil properties, productivity, fire regime, habitat structure, and species composition and their relative abundance [7,8]. Among them change in nutrient cycling is the most commonly reported example of ecosystem impacts of invasive species [7]. However, community structure (such as species richness and composition) changes long before the impact of invasive species on nutrient cycling is detected [8]. Decline in species richness due to invasion has been reported in several studies [1]. Similarly, change in species composition without significant change in species richness has been also reported [9]. The impact of invasion on community structure depends on trophic position of invader, taxonomic position of invader as well as the organism in the recipient community being considered, the ecosystem/habitat type, and disturbance regimes [1,10]. It is important to understand changes in species composition of the invaded community and growth dynamics of the invasive species itself over the time so that it would be possible to devise proper management strategies for the control of invasive plant species.

In Nepal, out of 179 naturalized plant species (Shrestha et al. 2017), *Parthenium* is arguably one of the most damaging invasive weeds in grassland [6,14]. It is also problematic in agroecosystems of Asia, Africa and Oceania [15,16]. In Nepal, *Parthenium* was first reported in 1967 [13]. Since then, the weed has been spreading continuosly and become widespread throughout Nepal upto 2000 m elevation due absence of a rigourous control program [6,14]. Although several environmental impacts of *Parthenium* are widely studied, the temporal growth pattern of this plant species has been merely studied [15–17]. Soil nutrients, habitat quality, seed dispersal and disturbances are some of the important factors that contribute to the growth of invasive species during one of the four stages of invasion process: introduction, existence as casual aliens, naturalization and spread [18]. Out of many factors, grazing is also considered as important that contributes in plant invasions [19] and is important to see how it helps invasive plant growth during certain time period.

*Parthenium* weed has been reported to reduce biodiversity in natural ecosystems including rangelands and protected areas [15]. However, only few studies examined plant species diversity in *Parthenium* invaded sites using appropriate experimental designs and there was lack of consistent pattern in the impacts of *Parthenium* on plant species diversity of the recipient community [20–22]. Nigatu et al. [21] reported a decline in species diversity (Shannon’s index) with increasing cover of *Parthenium* in grazing lands of Ethiopia. A similar decline in species diversity (Shannon’s index; aboveground as well as seedbank) in quadrats with high *Parthenium* density was reported in a grazing grassland of Queensland, Australia [20]. But, Timsina et al. [22] showed higher species richness (number of species per 1 m^2^ quadrat) in intermediately invaded quadrats than in the adjacent non-invaded quadrats of the grazing grasslands in Nepal. In addition to the lack of consistent patterns of the impact, the previous studies did not examine how species diversity in *Parthenium* invaded sites varies over time. It is highly likely that abundance of *Parthenium* weed as well as the species diversity of the invaded community change over time and understanding of this pattern is important for planning management of *Parthenium* weed. There are cases when abundance of invasive species declines (‘bust’ phase), even in absence of management interventions, after a period of high abundance (‘boom’ phase) due to processes mainly related to invasion and environmental changes [23]. To understand dynamics of *Parthenium* weed abundance and plant community structure in the invaded grazing grassland, we specifically addressed the following research questions: (1) How does growth of *Parthenium* (height, density, cover and soil seedbank density) vary between years in central Nepal? (2) How does soil seedbank of *Parthenium* change over a time period and vary according to soil depth? (3) How species richness and composition of different plant species change over certain time period in a grassland invaded by *Parthenium*? To answer above-mentioned questions, we collected the data on *Parthenium* density, cover, and height in different quadrats, associated plant species, grazing intensity, and soil seedbank of *Parthenium* from 2009 to 2013 in Hetauda, south-central Nepal.

## Materials and methods

### Study area

The study was undertaken at Lamsure Danda of Hetauda Municipality in Makawanpur district, south-central Nepal. The study site (27° 25’ N latitude, 85° 03’ E longitude, 500 m a.s.l.) lies in a dun valley of Siwalik region. The region has four distinct seasons: cold and dry winter (December-February), hot and dry summer (March-May), hot and humid monsoon (June-August) and moist and warm fall (September-November). Analysis of weather data between 1976 and 2005 revealed that annual precipitation was 2430 mm with monsoon rain contributing to about 82% [24]. Mean minimum and maximum temperatures were 16.3° and 29.2° C, respectively. *Shorea robusta* Gaertn. is the dominant tree species in forests of the region. The area lies in Tarai-Duar Savanna and Grassland eco-region of the world [25].

The study grassland is a property of government owned Hetauda Cement Factory but local communities have been using the grassland for grazing livestock and harvesting forage. The area was used for farming by local communities until 1977. In 1978, the area was purchased by Hetauda Cement Factory for mining soil and since then the site remained abandoned. This ‘secondary’ grassland has the highest species richness of the invasive alien plants among the natural vegetation in the region [26]. *Parthenium, Lantana camara* L., *Senna tora* (L.) Roxb., *Xanthium strumarium* L., *Spermacoce alata* Aubl. are the common invasive alien plants in this grassland while *Clerodendrum viscosum* Vent., *Imperata cylindrica* (L.) Beauv., *Chrysopogon aciculatus* (Retz.) Trin. are the common native species. The area was not subjected to any weed management. A few instances of slashing, uprooting and firing of *Parthenium* weed were observed but these activities were not common and less likely to have significant impacts on abundance of *Pathenium* weed in the study site.

### Parthenium survey and associated plant sampling

Sampling was carried out during August-September from 2009 to 2013. In the study area, three sites were selected that were about 150m apart with relatively high cover (>50%) of *Parthenium*. At each site, 10 square quadrats each of 1m^2^ with >50% cover of *Parthenium* were defined within big ca. 20m × 20m plot. Altogether, there were 30 quadrats sampled each year. Vascular plants growing in each quadrat, *Parthenium* growth (cover, density, and height of the tallest plant) were noted down. *Parthenium* cover was estimated visually in percentages, *Parthenium* density was counted in number (plant individuals within 1m^2^ quadrat) and height was measured in centimeter. Grazing intensity in each quadrat was also estimated in the scale of 0 (no sign of livestock grazing) to 3 (vegetation removal by livestock in >50% area of the quadrat). Each quadrat was marked by fixing a green colored polyvinyl chloride (PVC) pipe (length, 40 cm).

Associated plant species were recorded in each quadrat and specimens were collected for herbarium. Identification was carried out by comparing specimens deposited at National Herbarium and Plant Laboratories (KATH, Godawari, Lalitpur, Nepal) and referring ‘Plant Diversity of Eastern Nepal: Flora of Plains of Eastern Nepal’ [27]. Nomenclature was followed from Press et al. ([28] and The Plant List [29].

### Soil sampling and seedbank

Each quadrat was relocated in November with the help of fixed PVC pipe for soil sampling when most of the plants including *Parthenium* dispersed their seed. Soil samples were collected with the help of soil core sampler of ten centimeters diameter from the center of each quadrat after removal of ground vegetation and plant litter (but not seeds). Soil was collected in two steps: i) from surface up to 5cm depth, and ii) from 5cm up to 10cm depths. Altogether there were 60 soil samples collected from 30 quadrats each year. Soil samples were kept in plastic bag and brought to laboratory at Central Department of Botany, Tribhuvan University, Kathmandu for further analysis.

In each collected soil sample, seedling emergence method was used to assess the germinable soil seedbank (hereafter *Parthenium* seedbank) as it was difficult to extract seeds from the soil and count them [30]. Collected soil samples were kept into germination for two months in the greenhouse without any controlled environment to determine *Parthenium* seedbank. For this, earthen pots each of 15cm in diameter were taken and in each pot 2/3rd parts were filled with heat sterilized sand and then soil sample was spread over it, uniformly such that depth of the soil sample was less than 2cm. Sand and soil sample was separated by a layer of newspaper to prevent their mixing. All the pots were watered regularly with 100ml of water for each pot. They were observed regularly for newly emerging seedlings. Number of seedlings *Parthenium* that emerged in each pot were recorded weekly until the end of the experiment (i.e. two months). In each observation, seedlings were removed to avoid crowding.

### Data analysis

The analyses were carried out in three steps. First, we explored the determinants of *Parthenium* growth (cover, density and height). Here, we used *Parthenium* height, cover and density as response variables, and year, grazing and their interactions were used as explanatory variables. Second, to find out determinants of *Parthenium* seedbank, we used number of emerged seedlings as response variable and year, depth of soil, grazing, and their interactions as explanatory variables. Third, we assessed determinants of species richness of associated species. Here, number of different plant species growing in each quadrat (1 m^2^) was termed as species richness. In this analysis species richness was used as response variable, and explanatory variables used were year, grazing, *Parthenium* cover/density/height and their interactions.

All analyses were carried out by using the generalized linear mixed model (GLMM) with Poisson distribution using lme4 function in lmerTest package in R 3.3.2 [31]. We used Poisson distribution because data were normally distributed. In all tests, the effect of each quadrat was also taken into account by using them as random factor in above mentioned models. To derive the p-value, we first used drop1 function implemented in R and then again used Chi-square test [31]. All the figures were drawn by using STATISTICA 12 [32]. To find out variation in significant values from one another, we used Tukey’s post-hoc Test in Statistica 12.

Multivariate test was used to see the effect of *Parthenium* cover, density and height and also grazing on composition of associated plant species recorded in different quadrats. We used Redundancy Analysis (RDA) because of a short gradient length of 2.01 [33] by using Canoco 5.12 [34]. In RDA, model was built in the same way as it was done in GLMM. First, we tested significance of year and grazing intensity. If they were significant then the significant factor was taken as co-variate. In the final model, we tested the effect of year, grazing intensity (only significant variables), *Parthenium* cover, height, density and their interactions on species composition of associated plant species. The significance was tested by performing Monte Carlo permutation test (n=4999). To make the test possible, we added fictive species which was present in all the sampling quadrats. In RDA, we down weighted the rare species to reduce their effect on the results.

## Results

The correlations test showed that *Parthenium* cover was positively correlated with *Parthenium* density (p<0.001, r= 0.414) and plant height (p<0.001, r= 0.598). From the year 2009, *Parthenium* cover and height significantly decreased in 2010 but increased afterward (Figure 1a, b, Table 1). *Parthenium* density decreased from 2009 to 2010 and then the pattern was variable in remaining years (Figure 1c, Table 1). *Parthenium* density ranged from 15-703 stem m^−2^ whereas height ranged from 26-188cm. All these growth parameters of *Parthenium* decreased along with increasing grazing intensity. There was significant effect of interactions of year × grazing on *Parthenium* cover and density in our study area (Table 1).

**Table 1.** Relationship between *Parthenium* cover, density and height with year, grazing intensity and their interactions obtained from generalized linear mixed model (GLMM) test.

|  | Df | *Parthenium* cover |  | *Parthenium* density |  | *Parthenium* height |  |
| --- | --- | --- | --- | --- | --- | --- | --- |
|  |  | p-value | R<sup>2</sup> | p-value | R<sup>2</sup> | p-value | R<sup>2</sup> |
| Year | 1 | <0.001 | 0.474 | <0.001 | 0.754 | <0.001 | 0.437 |
| Grazing | 1 | <0.001 | 0.457 | <0.001 | 0.232 | <0.001 | 0.555 |
| Year × Grazing | 1 | 0.0023 | 0.069 | <0.001 | 0.013 | 0.2167 | - |

**Figure 1a.**
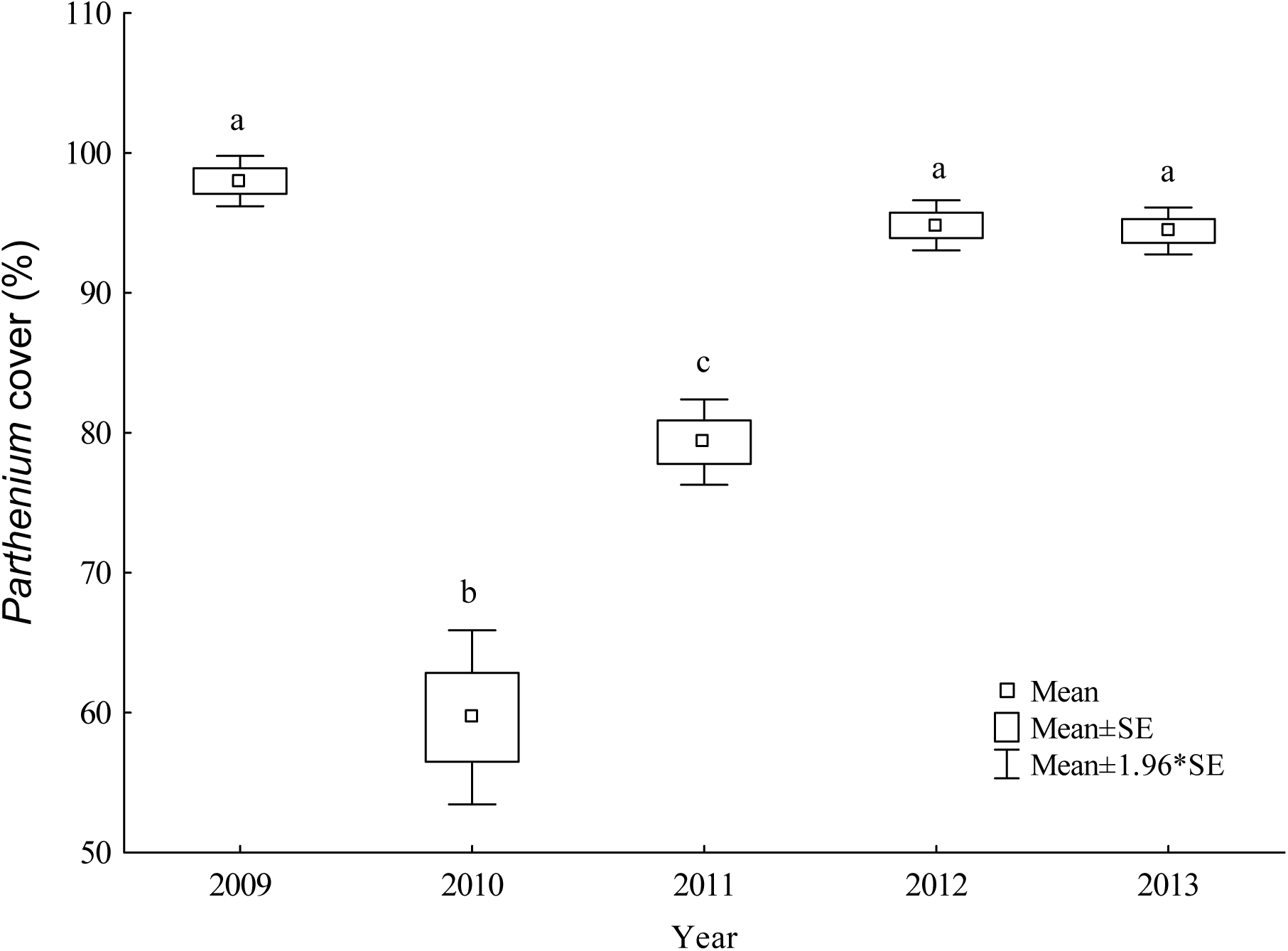
Variation of *Parthenium* cover over the study period.

**Figure 1b.**
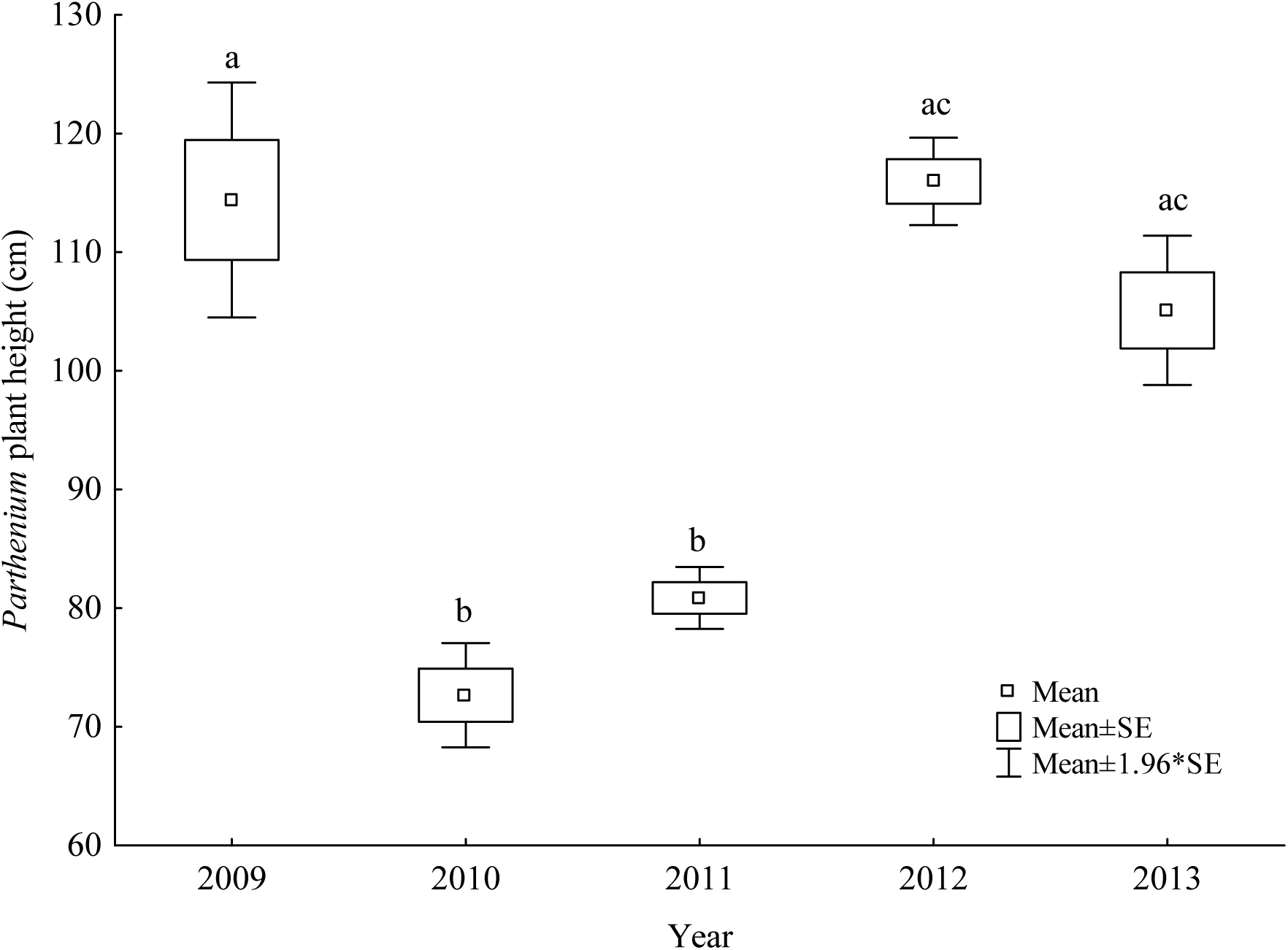
Variation of height of *Parthenium* over the study years.

**Figure 1c.**
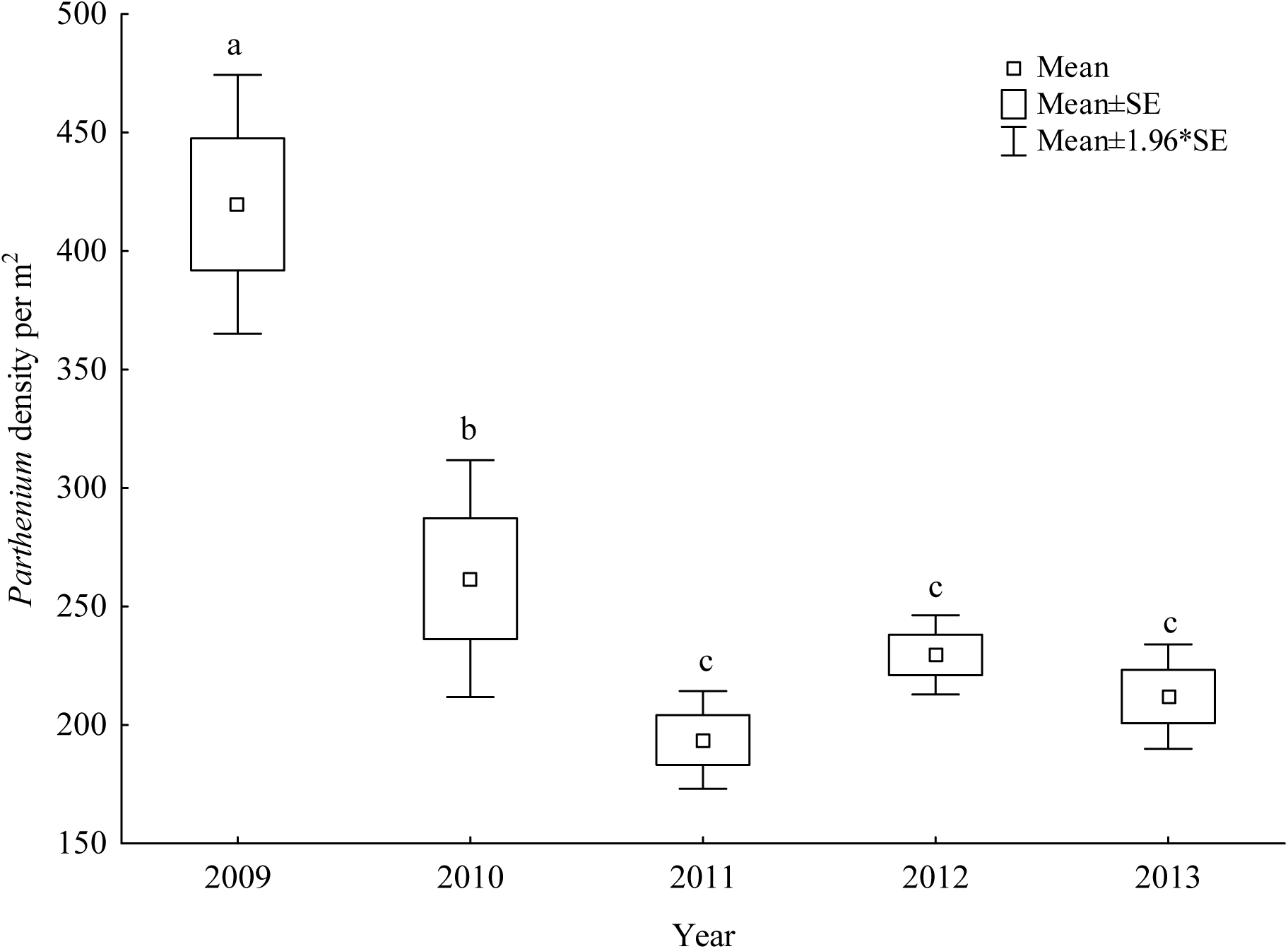
Variation of *Parthenium* density over the stud years.

Seedling emergence showed that *Parthenium* seedbank was eight times higher in shallow soil depth than the deep soil (Figure 2a). There were 127 to 42,675 seeds m^−2^ in shallow soil and no seeds to 16,815 seeds m^−2^ in deep soil. There was significant increase in *Parthenium* seedbank from 2009 to 2012 but a sharp decline in 2013 (Figure 2b). Though the seedbank was affected by year, soil depth, grazing intensity, and their interactions, the soil depth was the most important factor in maintaining seedbank (Table 2). Interactions of year × depth is the second most important factor in maintain soil seedbank whereas other significant factors have less contributions.

**Table 2.** Determinants of *Parthenium* soil seedbank obtained from generalized linear mixed model (GLMM) test.

|  | Df | p-value | R<sup>2</sup> |
| --- | --- | --- | --- |
| Year | 1 | <0.001 | 0.002 |
| Soil depth | 1 | <0.001 | 0.832 |
| Grazing | 1 | <0.001 | 0.002 |
| Year × soil depth | 1 | <0.001 | 0.114 |
| Year × grazing | 1 | <0.001 | 0.048 |
| Soil depth × grazing | 1 | <0.001 | 0.001 |
| Year × soil depth × grazing | 1 | <0.001 | 0.001 |

**Figure 2a.**
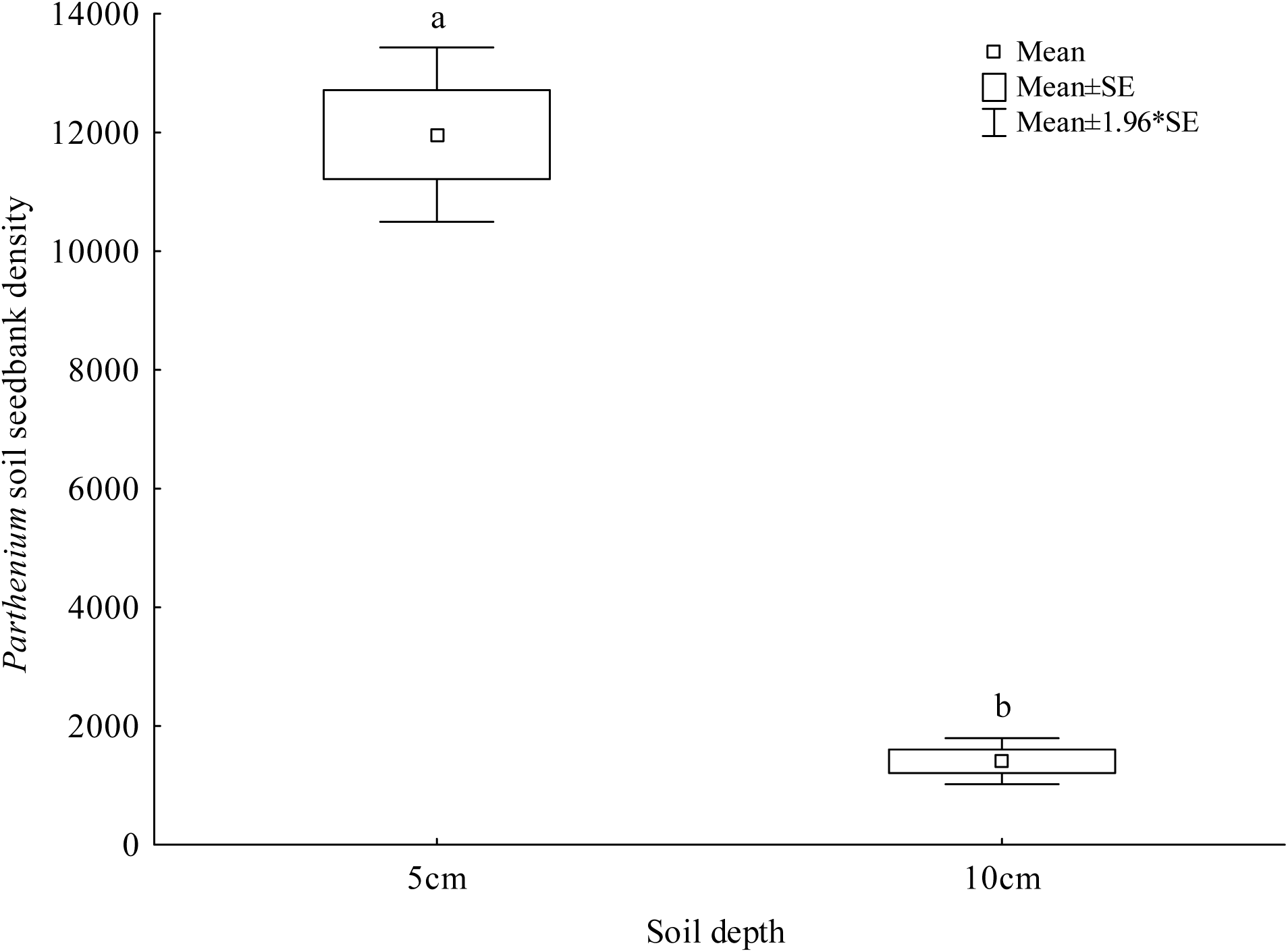
Change in *Parthenium* seed bank with soil depth.

**Figure 2b.**
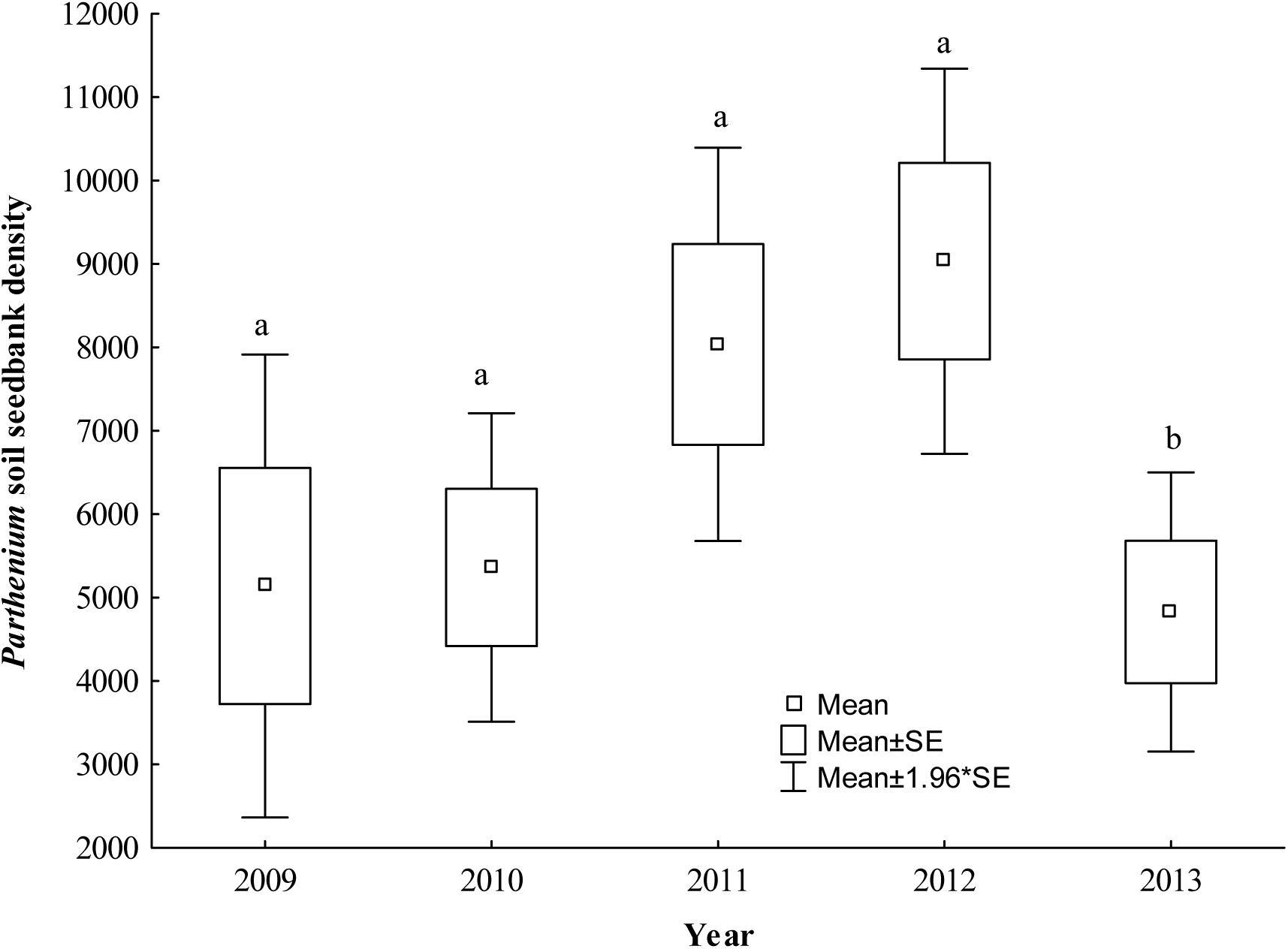
Change in *Parthenium* seed bank over the study years.

Out of all 87 plant species recorded in *Parthenium* invaded sites, 81 species (12 species identified up to genera level) belonging to 62 genera and 21 family were identified (Supplementary Table 1); 6 species could not be identified due to absence of reproductive parts in the specimens collected during field sampling. It was found that species richness of plant species significantly decreased until 2012 but slightly increased in 2013 (Table 3, Figure 3). Species richness of associated plant species decreased with increasing *Parthenium* height (Table 3, Figure 4). There was a significant effect of year × *Parthenium* density on the plant species richness (Table 3).

**Table 3.** Determinants of species richness of plant species in *Parthenium* invaded quadrats obtained from generalized linear mixed model (GLMM) test and determinants of species composition obtained from Redundancy Analysis (RDA) test.

|  | Species richness |  |  | Species composition |  |
| --- | --- | --- | --- | --- | --- |
|  | Df | p-value | R<sup>2</sup> | p-value | R<sup>2</sup> |
| Year | 1 | <0.001 | 0.627 | 0.434 | - |
| Grazing | 1 | 0.506 |  | 0.030 | 0.008 |
| *Parthenium* cover (cover) | 1 | 0.055 | - | 0.002 | 0.027 |
| *Parthenium* height (height) | 1 | 0.001 | 0.146 | 0.264 | - |
| *Parthenium* density | 1 | 0.218 | - | 0.002 | 0.019 |
| Year × grazing | 1 | 0.121 | - | 0.006 | 0.024 |
| Year × cover | 1 | 0.252 | - | <0.001 | 0.013 |
| Year × height | 1 | 0.95 | - | 0.022 | 0.013 |
| Year × density | 1 | 0.022 | 0.089 | 0.006 | 0.024 |
| Grazing × cover | 1 | 0.133 | - | 0.006 | 0.045 |
| Grazing × height | 1 | 0.497 | - | 0.009 | 0.033 |
| Grazing × density | 1 | 0.118 | - | 0.032 | 0.032 |
| Cover × height | 1 | 0.473 | - | 0.013 | 0.034 |
| Cover × density | 1 | 0.886 | - | 0.043 | 0.033 |
| Height × density | 1 | 0.738 | - | 0.046 | 0.022 |

**Figure 3.**
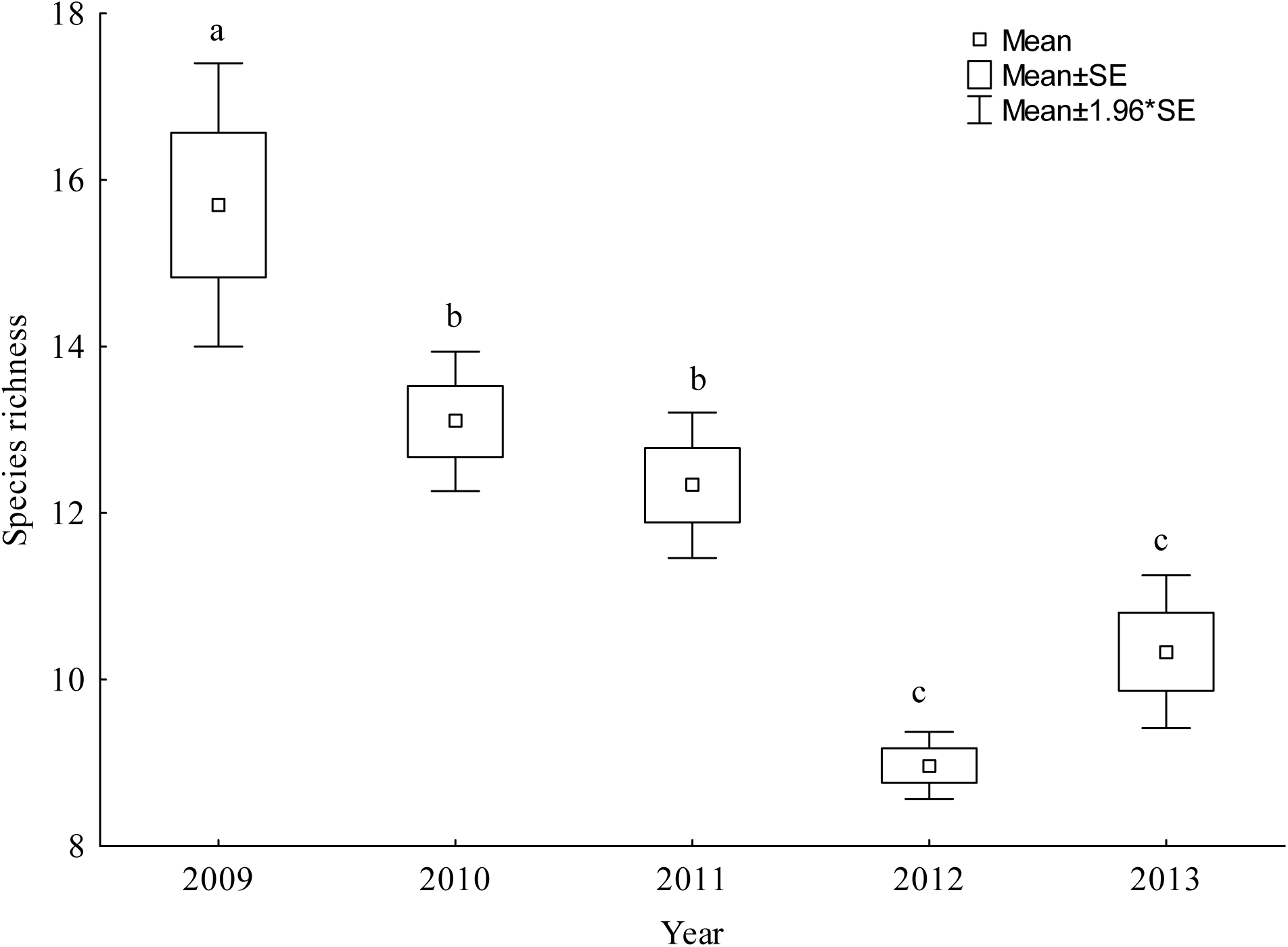
Variation of species richness of the associated plant species in *Parthenium* invaded quadrats over the study years.

**Figure 4.**
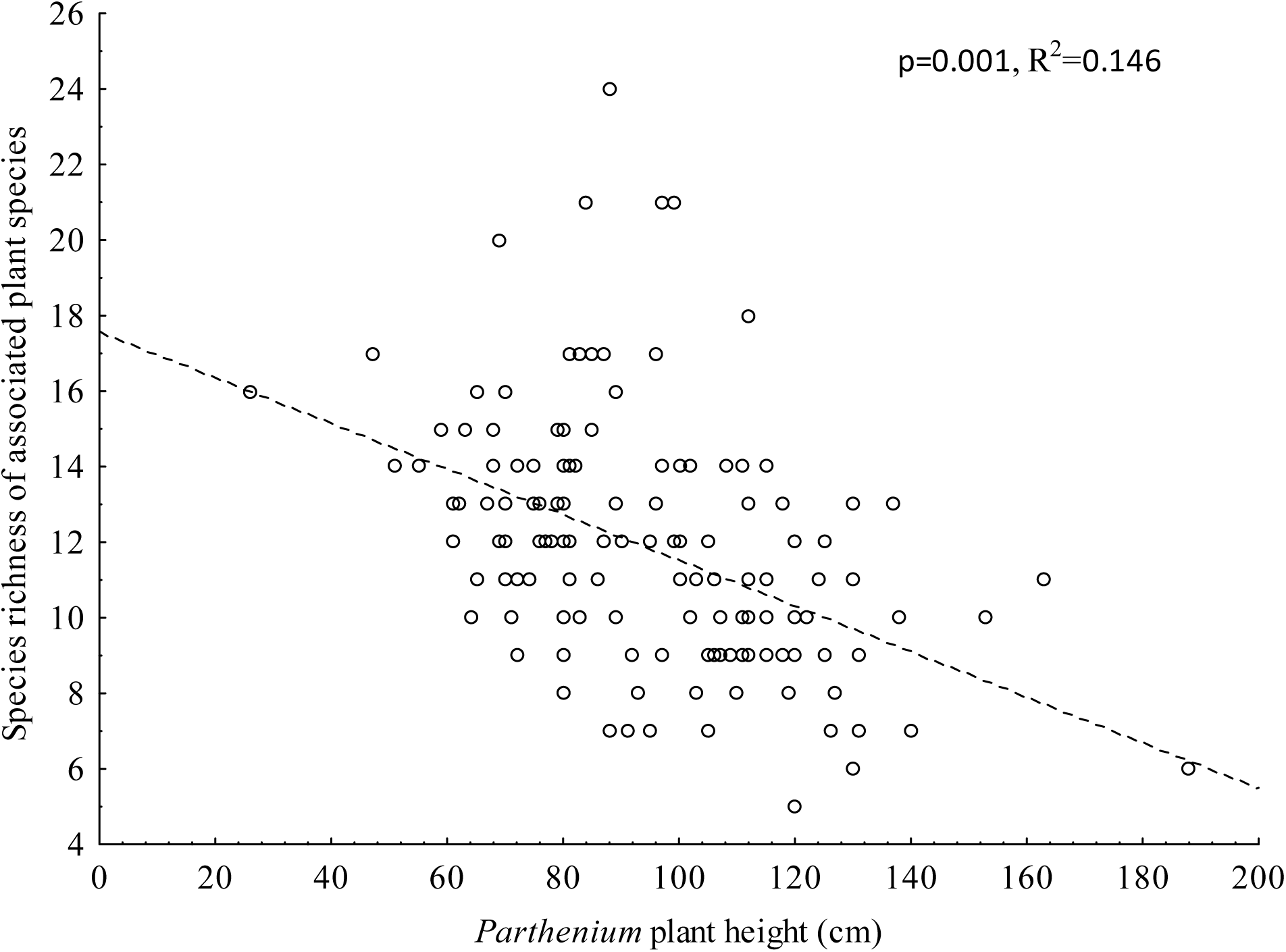
Relationship between species richness and *Parthenium* plant height.

The multivariate analysis of species composition of associated species was significantly affected by grazing intensity, *Parthenium* density and cover (p=0.039). Similarly, all two ways interactions of different factors affected species composition (Table 3).

Associated plant species that were found in quadrats with high grazing intensity and low *Parthenium* cover were *Sida rhombifolia* L., *Oxalis corniculata* L., *Crotalaria prostrata* Rottler ex Willd., *Phyllanthus urinaria* L. and *Lindernia parviflora* (Roxb). Haines. Similarly, associated species which grew in areas with high *Parthenium* density were none but in areas with medium density of *Parthenium* were *Oplismenus burmannii* (Retz.) P. Beauv, *Setaria glauca* (L.) P.Beauv and *Ageratum conyzoides* L.. *Axonopus compressus* (Sw.) P. Beauv., *Ophioglossum* sp., *Leucas indica* (L.) Sm. and *Desmodium triflorum* (L.) DC were found in quadrats where there was high *Parthenium* cover (Table 3, Figure 5).

**Figure 5.**
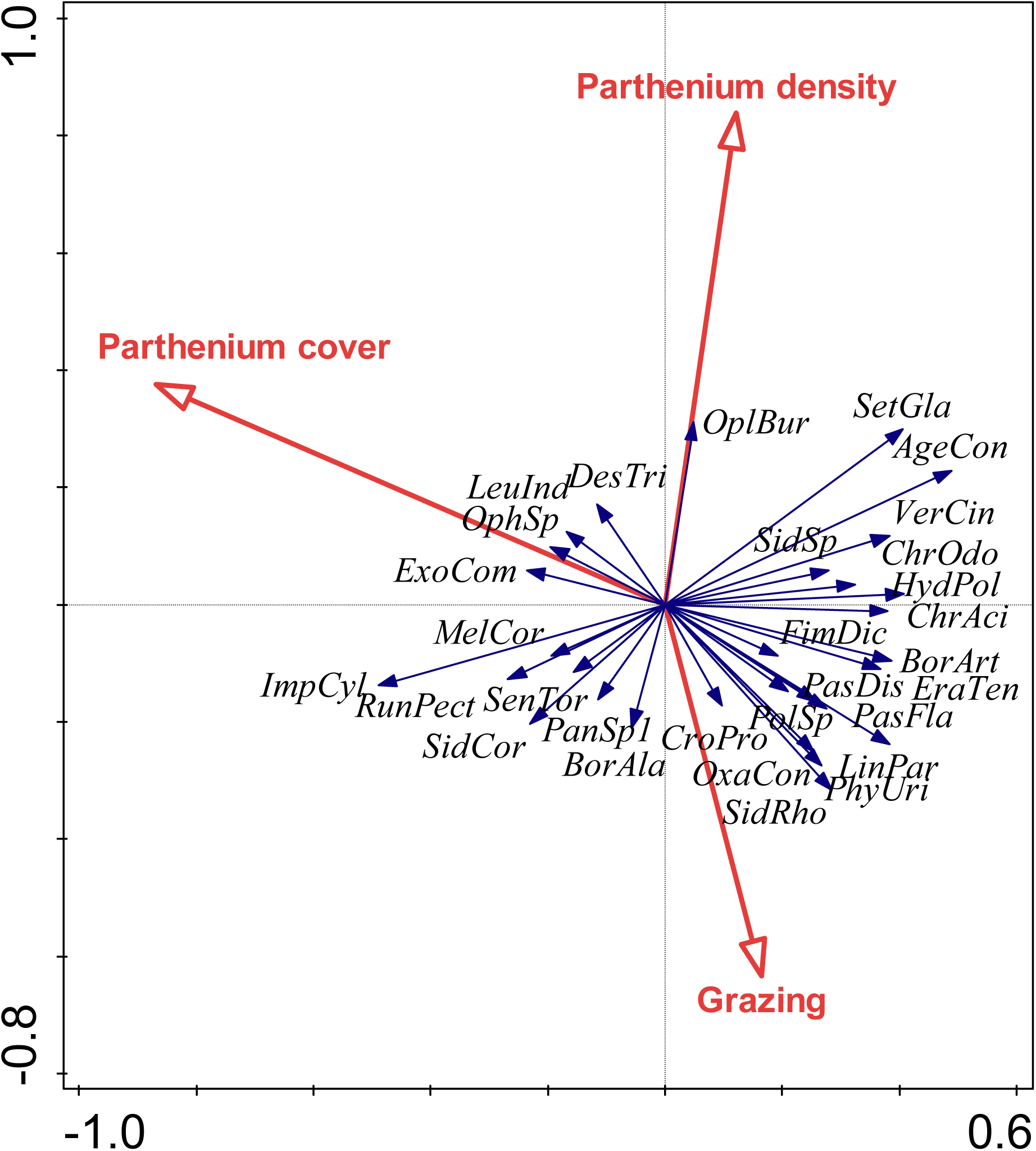
Relationship between *Parthenium* associated plant species with grazing and *Parthenium* plant density. The 1^st^canonical axis explained 6.12% and the 2^nd^ 2.56% of the total variation in the data set. Abbreviations for plant species are given in Supplementary Table 1.

## Discussion

Long-term planning for effective management of invasive alien plants require an understanding of how species composition of the invaded community and abundance of the invasive species itself change over time. Using data collected for five years in a *Parthenium* invaded grazing grassland of south-central Nepal, we showed that despite annual variation the abundance of *Parthenium* has in general increased over time with a significant shift in species composition. The results indicate that the persistence of many plant species will be threatened seriously with direct negative impacts on biodiversity, ecosystem processes and livestock farming if management of *Parthenium* is not initiated immediately.

The invasion of *Parthenium* has increased over the time in our study site, and a similar situation could be expected in other regions invaded by *Parthenium* in Nepal. This fact is illustrated by our observations showing increase of *Parthenium* cover and height of the plant individuals from 2010-2013. Rate of increase of invasive plant with time is also found in different studies in Nepal and around the world [6,15,35,36]. The decrease in *Parthenium* cover, density and height from the year 2009 to 2010 might be due to the herbivore damage caused by a biological control agent *Zygogramma bicolorata* (Coleoptera: Chrysomelidae) that was first reported from Hetauda in Nepal in 2009 [37]. This beetle is considered an effective biological control agent that feed on leaves of *Parthenium* and significantly reduce growth, vigor and seed production of this plant [5]. However, there may be year-to-year variation in the damage caused by *Z. bicolorata* on *Parthenium* weed due to variation in weather conditions [38].

The positive correlation between cover with density and height of *Parthenium* in our study shows that either cover or density could be used to represent one another. *Parthenium* density in our study (15-703 stem m^−2^) is very high compared to data available from Pakistan [39]. However, previous study by Karki [40] in the same region showed a mean density of 118 stem m^−2^ showing a sharp increase in abundance of *Parthenium* over a time period. The plant height in our study (26-188cm) was less than in highly invaded areas in Australia where they are up to 2.5m in height although most plants do not exceed 1.5m [41]. Mean height estimated from different parts of Nepal is 128cm, with maximum value reaching up to 240cm [14]. It appears that the maximum height reported from Nepal and Australia are similar but it was lower in the present study site. Either edaphic or climatic factors might have prevented the individuals of the present study site to grow taller. In addition, negative impacts of defoliation by the biological control agent could not be ruled out but this was not evaluated in the present study.

*Parthenium* is normally avoided by cattle [6] but grazing intensity varied with the abundance of *Parthenium*. Low grazing intensity in highly invaded quadrats (i.e. with high abundance of *Parthenium*) suggests low visitation by cattle, probably due to low availability of forage in such quadrats. Reduction of native plant biomass nearly to 1/3^rd^ has been reported due to dense growth of *Parthenium* in three dun valleys of Nepal including Hetauda [14].

Prevalence of more seeds in shallow top soil compared to deep soil is an well-established phenomenon [30,42]. These seeds remain viable for a very long time period and eradication of *Parthenium* is difficult once it starts growing in some places. Adkins and Shabbir [15] have found that more than 70% of seeds below 5cm remain viable for 2 year, with a half-life of 7 years. Similarly, White ([43] found seeds to be viable for 4 to 6 years and other studies [44,45] also reported that buried seeds remain viable for much longer period than seeds on the soil surface. The impact of interactions of year × depth means that there is variability of seed bank according to the soil depth and is variable in different years. This is probably because of the plant growth and seed production that vary from one year to next.

There was high number of seed production (up to 42675 seeds m^−2^ accumulated in shallow soil and 16815 seeds m^−2^ in deep soil) by *Parthenium* in our study site. This is similar to other studies [20,45] but far less than the report by Joshi (1991) in India, who reported about 200000 seeds m^−2^. The increase in seedbank size could be attributed to the addition of seed to the persistent soil seed bank of *Parthenium* [15,43–45]. Increasing soil seedbank from 2009-2012 was alarming, which suggests the need of immediate management to prevent further degradation of the grassland due to *Parthenium* invasion. Sharp drop in seed bank from 2012 to 2013 might be due to low production of seeds as the year 2013 was exceptionally dry due to low precipitation [47].

In Hetuda, we found high number (n=87) of plant species in *Parthenium* invaded sites and many of them are palatable meaning that the site is rich in plant community. This number in our study is higher compared to other studies in Nepal [22,48], 72 species in Ethiopia [21], 30 associated species in Pakistan [49] and in different part of the world [15]. The sharp decrease in plant species richness from 2009 to 2012 have shown that invasion of *Parthenium* is ever increasing with serious adverse effect on many native species. This adversity will in near future may lead to severe degradation of habitat with loss of biodiversity [22] and brings about changes in existing ecosystem in larger scale [6,15]. A significant effect of year × *Parthenium* density on the plant species richness means that the density of *Parthenium* is variable in different years and its’ growth probably depends on favorable temperature or precipitation and disturbance in a year.

Effects of different environmental factors on species composition of the *Parthenium* invaded community is in line with other studies [6,15,21,22]. Each environmental factor acts differently and thus result in differences in associated plant species composition in a given space. Taxonomic identity of plant species growing in *Parthenium* invaded sites differ from place to place as shown in Nepal [22] and elsewhere in the world [15,21]. Our multivariate analysis shows that high number of associated plant species do not prefer *Parthenium* invaded quadrats resulting in variation in species composition of associated plant species. However, there was no effect of time period on associated plant composition meaning that despite *Parthenium* invasion most of the plant species still can persist at quadrats where the abundance of *Parthenium* is low. This is also in agreement with our previous report that plant species richness can be higher in plots that are intermediately invaded by *Parthenium* than in the heavily invaded, or non-invaded quadrats [22].

The results show that *Parthenium* is ever increasing in our study site at Hetauda. In addition to habitat degradation and changes in community structure, the increase in invasion severity may in future cause serious problems in health for both livestock and human due to its toxic nature [20,21]. A wide range of management plans have been reported recently by Adkins and Shabbir ([15]. However, depending on human and economic resources available, the locally most appropriate and cost-effective methods should be used for *Parthenium* management. Further, as there are many important species in *Parthenium* invaded sites a proper competition experiment should be conducted to identify the vulnerable and competitive species in the communities. The competitive species can potentially be used as suppressive plants as a part of integrated management of *Parthenium* weed [50].

## Conclusion

Present multi-year study in a grazing grassland at Hetauda in south-central Nepal has provided in-depth knowledge on dynamics of *Parthenium* weed invasion. It has been found that the weed is a serious threat to the associated plant species that are palatable and important to maintain existing ecosystem. Variation of species richness of plant species over 5-year time period and also increasing size of soil seedbank shows that *Parthenium* is currently a potential threat to biodiversity that is also increasing with time due to accumulation of seeds in seed bank. Thus, awareness campaign, formulation of proper management plans and their implementation is needed to avoid serious threats posed by *Parthenium* weed invasion in different parts of Nepal.

## Author Contribution

BBS and MBR conceived and designed the study. JKC, BBS, SG and MBR collected data. MBR analyzed data. MBR and BBS wrote the manuscript and all other authors commented and approved the final version of manuscript.

## Acknowledgement

This study was supported by Nepal Academy of Science and Technology (NAST) for data collection of 2009 and 2010 for all except MBR. For MBR, the study was supported by the Czech Science Foundation (project no. 17-10280S) and partly by institutional support RVO 67985939. We are thankful to Binu Timsina for arranging data in proper format for analysis. We are thankful to Hetauda Cement Factory for allowing us to carry out our field surveys and also locals who highly cooperated during our entire study period. Ambika Poudel helped in data collection during 2009 and 2010. We also thank the Central Department of Botany, Tribhuvan University, Nepal for allowing us to use greenhouse facility.

## Supplementary File

**Supplementary Table 1.**
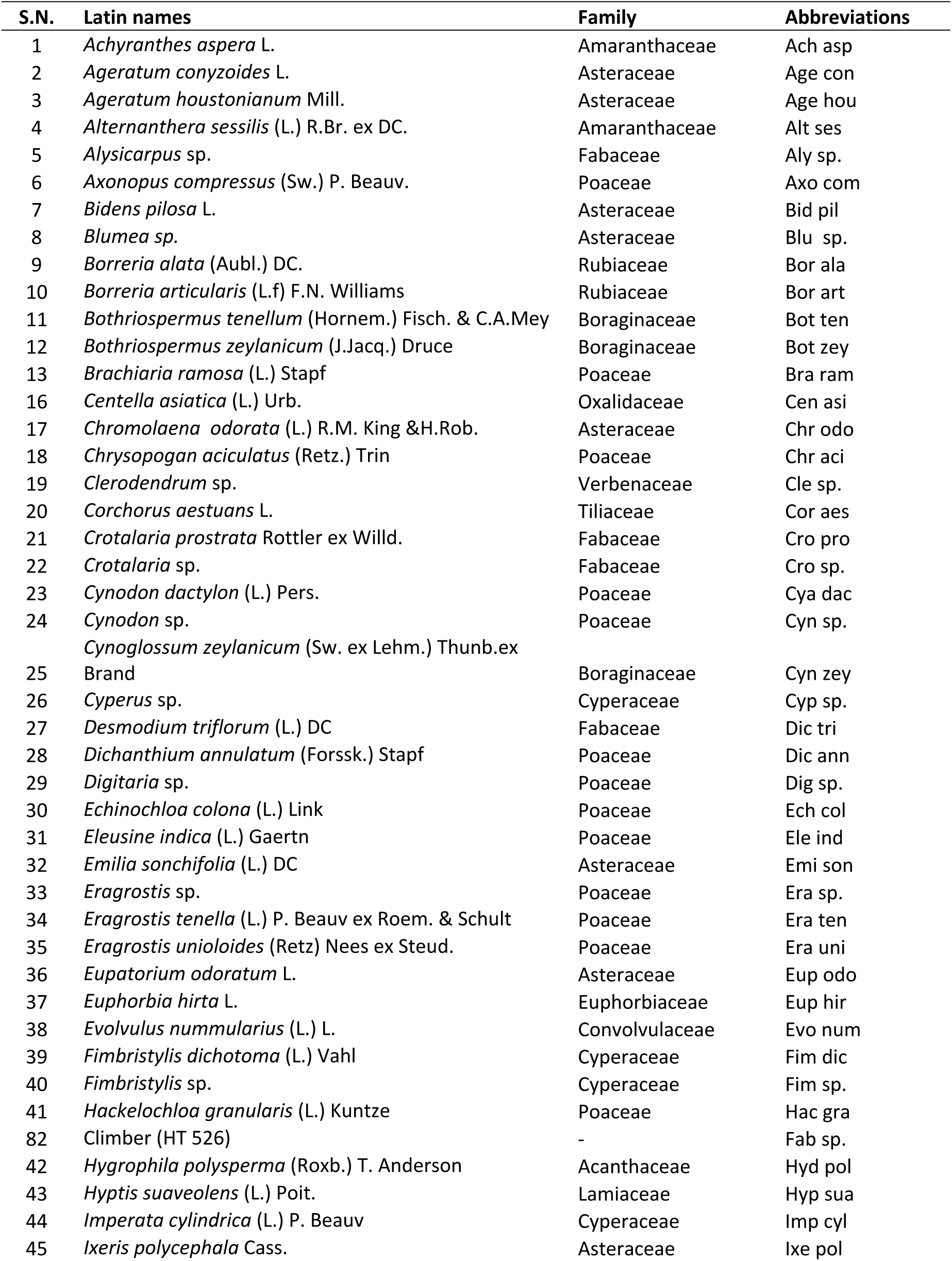
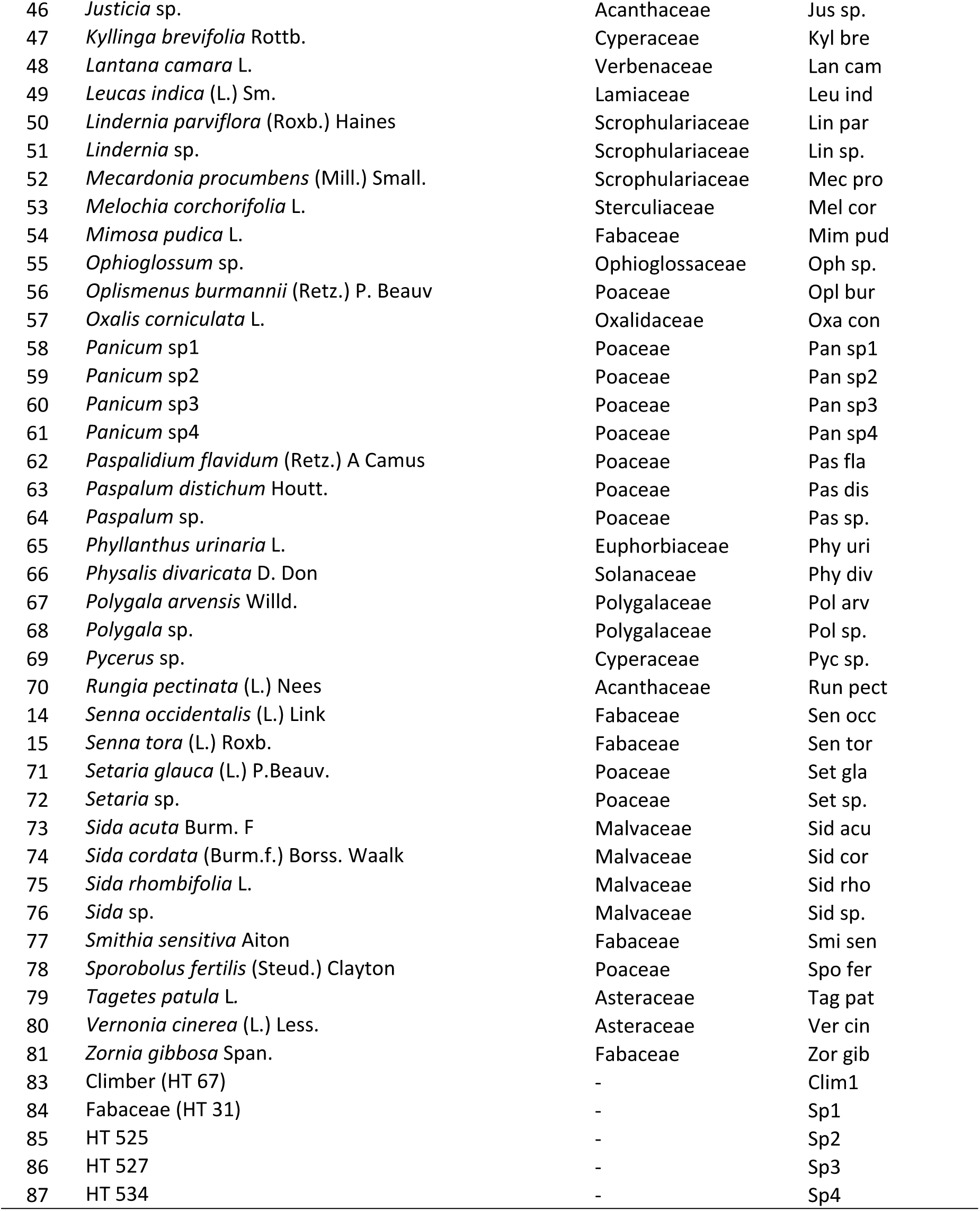
List of plant species recorded in *Parthenium* invaded quadrats.

